# Thalamocortical orchestration of human theory of mind

**DOI:** 10.64898/2026.07.28.741302

**Authors:** Runhao Lu, Emily Davidson, Jonah Miller-Smith, Dian Lyu, Zoe Lusk, Prantik Kundu, Julien Doyon, Xiaoqian J. Chai, Boris C. Bernhardt, Gary R. Turner, R. Nathan Spreng

**Affiliations:** Montreal Neurological Institute, Department of Neurology and Neurosurgery, McGill University, Montreal, Quebec, Canada; Laboratory of Behavioral and Cognitive Neuroscience, Department of Neurology and Neurological Sciences, Stanford University School of Medicine, Stanford, CA, USA; MOE Frontier Science Center for Brain Science and Brain-Machine Integration (BBMI), Zhejiang University, Zhejiang, China; Department of Neurosurgery, School of Medicine, the Second Affiliated Hospital of Zhejiang University, Zhejiang, China; Ben L. Silberstein Institute for Brain Health, Wayne State University, Detroit, MI, USA; Icahn School of Medicine at Mount Sinai, New York, NY, USA; McConnell Brain Imaging Centre, McGill University, Montreal, Quebec, Canada; Department of Psychology, York University, Toronto, ON, Canada

**Keywords:** Mentalizing, Thalamus, Partly Cloudy, Naturalistic, Default Mode Network (DMN), sEEG, 7T fMRI

## Abstract

Reasoning about others’ thoughts or beliefs is central to human social behavior. This ability, known as theory of mind (ToM), has been primarily attributed to cortical regions of the default network (DN). However, whether and how subcortical structures, particularly the thalamus, contribute to this high-level computation remains unknown. Here, we investigated human thalamocortical dynamics during a naturalistic ToM movie watching condition, taking a rare dual-modality approach by combining high-field 7T fMRI and intracranial stereoelectroencephalography (sEEG). Across both modalities, ToM events reliably activated the DN, whereas the thalamus lacked canonical local activation. Despite this absence of local activation, the thalamus shared ToM-related representational structure and exhibited enhanced bidirectional interactions with the DN during mentalizing across methods. Crucially, sEEG revealed that the thalamus coordinated DN activity via cross-frequency phase-amplitude coupling (PAC), whereby thalamic low-frequency phase unidirectionally modulated DN high-frequency activity. Furthermore, the strength of thalamic-DN PAC predicted both the activity magnitude and the representational quality of ToM-related information within the dorsomedial DN subsystem. Together, these findings identify the human thalamus as a regulatory hub that gates DN computations during ToM without exhibiting observable localized activity, revealing a previously unrecognized mechanism by which thalamic dynamics coordinate high-level social cognition in humans.

## Introduction

The ability to infer the beliefs, intentions, and mental states of others, termed “theory of mind” (ToM) or “mentalizing”, is a central component of human social cognition and supports flexible social interaction across diverse contexts ^1–6^. Utilizing controlled laboratory tasks ^7^ and naturalistic paradigms ^8,9^, functional neuroimaging and intracranial recordings have consistently localized ToM to a distributed set of cortical regions, including the temporoparietal junction, posteromedial cortex, and medial prefrontal cortex ^2,6,9–13^, that largely align with the core and dorsomedial subsystems of the default network (DN) ^10,14,15^.

However, this corticocentric framework is increasingly challenged by evidence that subcortical structures, particularly the thalamus, play a critical role in coordinating large-scale cortical computation. Moving beyond its classical characterization as a passive sensory relay, the thalamus is recognized as an essential coordinator of distributed cortical computation ^16–19^. This coordinating capacity is anchored in the dense, reciprocal projections between higher-order thalamic subdivisions (such as the mediodorsal and anterior groups) and the prefrontal regions that comprise the DN ^20–27^. Such circuits are crucial for shaping cortical dynamics ^28,29^, guiding attentional control ^17,30–32^, and enabling cognitive flexibility ^33^. In these contexts, the thalamus frequently acts as a regulatory hub that gates cortical information processing and maintains cortical representations ^20,33^, with cross-frequency phase-amplitude coupling (PAC) between thalamic and cortical regions serving as a putative key mechanism ^34–37^. More recently, evidence from rodent models has extended these thalamic functions into the social domain, demonstrating that the thalamus and its connectivity to medial frontal or posterior parietal cortex, key homologs of the human DN regions, actively encode social information and coordinate social memory ^38–40^.

Despite these pivotal insights, it remains largely unknown whether and how the human thalamus modulates large-scale cortical networks responsible for complex social cognition like ToM. Here we investigated the human thalamocortical dynamics underlying naturalistic ToM using a dual-modality approach, integrating high-field 7T multi-echo fMRI from 37 healthy adults, and stereoelectroencephalography (sEEG) from eight presurgical epilepsy patients with implanted electrodes (comprising 108 DN and 95 thalamic sites). Participants across both cohorts viewed the animated short film, “Partly Cloudy” ^41^, a well-validated movie containing temporally precise and annotated ToM events ^8,9,42^. We organized our analyses around three questions concerning the roles of thalamic-DN interaction in human naturalistic mentalizing. First, we asked whether the thalamus shows stronger localized activity during naturalistic ToM, comparable to the canonical activation expected in cortical DN regions. Second, given the extensive anatomical and functional connectivity between the thalamus and association cortex, we tested whether the thalamus carries ToM-relevant information and strengthens its communication with the DN during mentalizing. Third, based on the potential role of thalamus as a regulatory hub for cortical computation, we tested whether thalamic low-frequency dynamics rhythmically regulate DN activity during ToM via PAC, and whether this coupling supports cortical ToM processing by predicting the magnitude and representational quality of DN responses. We found that the human thalamus contributes to naturalistic ToM primarily through shared representational structure, enhanced thalamocortical communication, and rhythmic regulation of DN computations, rather than through stronger localized activity. These findings position the human thalamus as a background coordinator of cortical mentalizing, shaping foreground DN computations through rhythmic thalamocortical modulation.

## Results

To investigate how thalamic-DN interactions support ToM under naturalistic conditions, we combined a movie-watching paradigm with two complementary high-resolution recording modalities: high-field 7T fMRI and intracranial sEEG.

Participants viewed *Partly Cloudy* ^41^, a well-established animated film paradigm that reliably elicits spontaneous mentalizing. Prior work has identified a set of temporally precise ToM events embedded within the narrative ^8,9,42^, enabling event-related analyses of social inference. We leveraged these annotations to isolate seven ToM epochs throughout the movie (**Fig. 1a**; see **Supplementary Table S1** for details).

**Figure 1.**
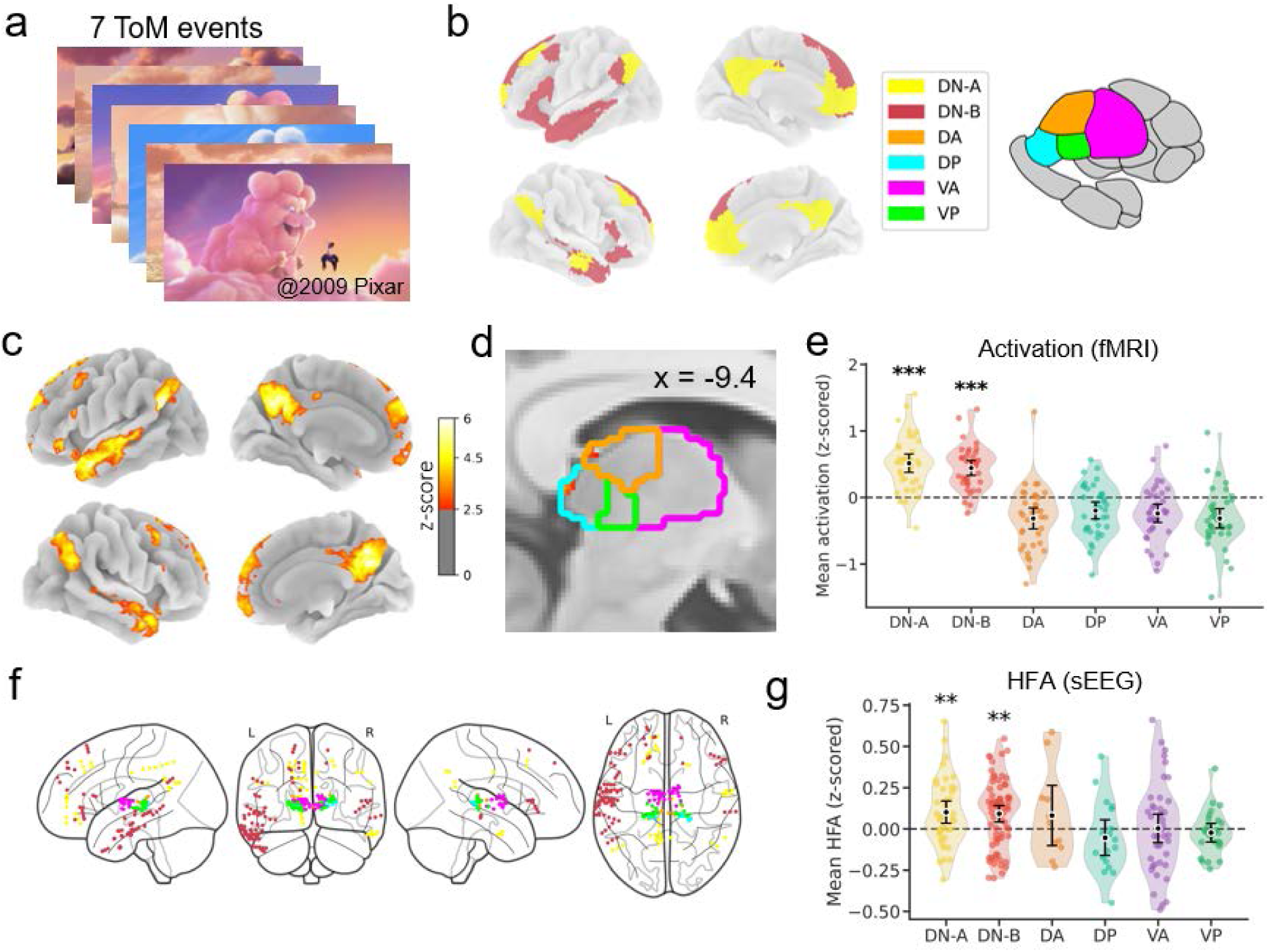
Naturalistic ToM paradigm, ROI definitions, and activation results of sEEG and fMRI. **(a)** Experimental paradigm using the animated short film *Partly Cloudy*. Seven temporally precise, annotated ToM events (defined in ^8^) were used for event-related analyses of naturalistic mentalizing. **(b)** Definition of cortical and subcortical regions of interest (ROIs). Cortical ROIs include the core (DN-A) and dorsomedial (DN-B) subsystems of the default network (defined by Schaefer 400 with Yeo 17 network parcellation ^43,44^). Thalamic subregions include dorsal anterior (DA), dorsal posterior (DP), ventral anterior (VA), and ventral posterior (VP) subregions defined by Melbourne Subcortex Atlas ^45^. **(c)** Group-level whole-brain 7T fMRI activation maps (n = 37) during ToM events (ToM > implicit baseline; FDR-corrected *p* < 0.05). **(d)** Group-level thalamic 7T fMRI activation maps during ToM (ToM > implicit baseline; FDR-corrected *p* < 0.05 within the thalamus). **(e)** ROI-based analysis of mean fMRI activation levels. Each dot represents an individual subject. Error bars represent 95% confidence intervals. **(f)** Anatomical distribution of sEEG recording sites across 8 patients (108 DN sites and 95 thalamic sites), color-coded by ROI. **(g)** ToM-related high-frequency activity (HFA; 70–170 Hz) during ToM events relative to the implicit movie baseline. Each dot represents an individual recording site within the respective ROI. ** *p* < 0.01, *** *p* < 0.001 (FDR-corrected).

Using a dual-modality approach combining 7T fMRI and sEEG, we extracted ToM-related signals from *a priori* defined regions of interest (ROIs) (**Fig. 1b**). Cortical ROIs focused on the core (DN-A) and dorsomedial (DN-B) subsystems of the DN, defined via the Schaefer 400-region parcellation ^43^ with the Yeo 17-network framework ^44^. Thalamic ROIs comprised four subregions delineated by the Melbourne Subcortex Atlas ^45^: dorsal anterior (DA), dorsal posterior (DP), ventral anterior (VA), and ventral posterior (VP). Our dataset comprised multi-echo fMRI data from 37 healthy young adults and parallel sEEG recordings from eight pre-surgical epilepsy patients, yielding a total of 108 DN sites (DN-A: 39, DN-B: 69) and 95 thalamic sites (DA: 11, DP: 18, VA: 41, VP: 25) (**Fig. 1f**). The 7T multi-echo fMRI acquisition yielded robust temporal signal-to-noise ratio (tSNR) across both the distributed cortical networks and all evaluated thalamic subregions (**Fig. S1**), ensuring reliable estimation of ToM-related signals in both cortical and subcortical ROIs.

### Cortical DN but not thalamus exhibits robust activation during naturalistic ToM

To determine a high-fidelity activation map of brain regions involved in naturalistic ToM, we first conducted 7T fMRI analyses in healthy adults. Whole-brain general linear model (GLM) analyses revealed widespread activation across DN regions during ToM events relative to the remainder of the film, encompassing both DN-A and DN-B subsystems (**Fig. 1c**). Marginal activation was observed within the thalamus, even under targeted small-volume correction, with minimal clusters in the dorsal subdivision (**Fig. 1d**). ROI-based analyses confirmed this pattern: DN-A and DN-B exhibited significant positive activation, whereas all thalamic subregions showed no significant ToM-related activation (**Fig. 1e**; FDR-corrected *p* < 0.05).

We next examined the pattern of brain activity for ToM using intracranial sEEG recordings. sEEG provides spatially focal measurements of local neural population activity from implanted sites, offering a direct electrophysiological test of whether ToM events elicit localized neural responses in these regions at a faster temporal scale. We quantified high-frequency activity (HFA; 70–170 Hz) as a proxy for local neural processing ^46,47^. During ToM events, both DN-A and DN-B exhibited significant increases in HFA relative to the implicit movie baseline (**Fig. 1g**; FDR-corrected *p* < 0.05). In contrast, none of the thalamic subregions showed significant HFA increases, indicating a lack of ToM-related activation across thalamic sites.

Together, these convergent findings across hemodynamic and electrophysiological modalities reveal a pronounced functional divergence in activation patterns within the thalamocortical system during naturalistic ToM. While the DN shows robust ToM-related activation, the thalamus does not exhibit measurable local activation, indicating that its contribution to social cognition is not reflected in conventional activation-based metrics.

### Thalamus shares ToM-related representations with the DN

The human thalamus is central to coordinated cognition, supporting distributed cortical computations through its interactions with association networks. Despite the absence of overt activation, we further probed hypothesized roles for the thalamus in ToM-related information sharing, thalamocortical communication, and rhythmic regulation of DN activity. First, we quantified the representational structure of neural activity during ToM using representational similarity analysis (RSA) in both 7T fMRI and sEEG (see Methods). This approach captures the temporal geometry of activity patterns within each region, allowing us to assess whether different regions share similar representational structure during mentalizing. We predicted that the thalamus would carry information related to the geometry of mental inference even in the absence of overt activity.

We first investigated this representational similarity in the fMRI hemodynamic signals using a leave-one-out (LOO) inter-subject RSA (IS-RSA) framework. This approach isolates stimulus-locked, shared representational structure across participants while mitigating the influence of subject-specific noise. We constructed TR-by-TR representational similarity matrices (RSMs) based on voxel-wise activity patterns during ToM events (**Fig. 2a**, top). To resolve the precise spatial topography of these shared representations, we first implemented a volumetric searchlight approach across the whole thalamus (see Methods). The resulting representational coupling maps revealed a highly organized topographical gradient within the thalamus during mentalizing (**Fig. 2b**). Specifically, strong positive representational alignment with both DN-A and DN-B was robustly concentrated in the dorsal thalamus. To quantitatively characterize these visually apparent spatial gradients, we performed a spatial binning analysis by parcellating the thalamus geometrically into three equal-volume bins along both the anterior-posterior (A-P) and dorsal-ventral (D-V) axes (**Fig. 2c**). This fine-grained quantification revealed structured topographical profiles that were consistent across both DN-A and DN-B subsystems, highlighting critical axis-specific effects. Along the A-P axis, a significant positive representational alignment against zero was selectively present in the posterior bin for the DN-A network (FDR-corrected *p* < 0.05). Crucially, along the D-V axis, both DN-A and DN-B exhibited clear spatial transitions, where representational coupling was significantly stronger in both the middle and dorsal bins relative to the ventral segment (FDR-corrected *p* < 0.05). Furthermore, when testing absolute coupling against zero, only the middle and dorsal bins within the DN-A pathway reached significance (FDR-corrected *p* < 0.05), indicating that the representational alignment with higher-order thalamic areas may be particularly prominent for the core DN subsystem in fMRI.

**Figure 2.**
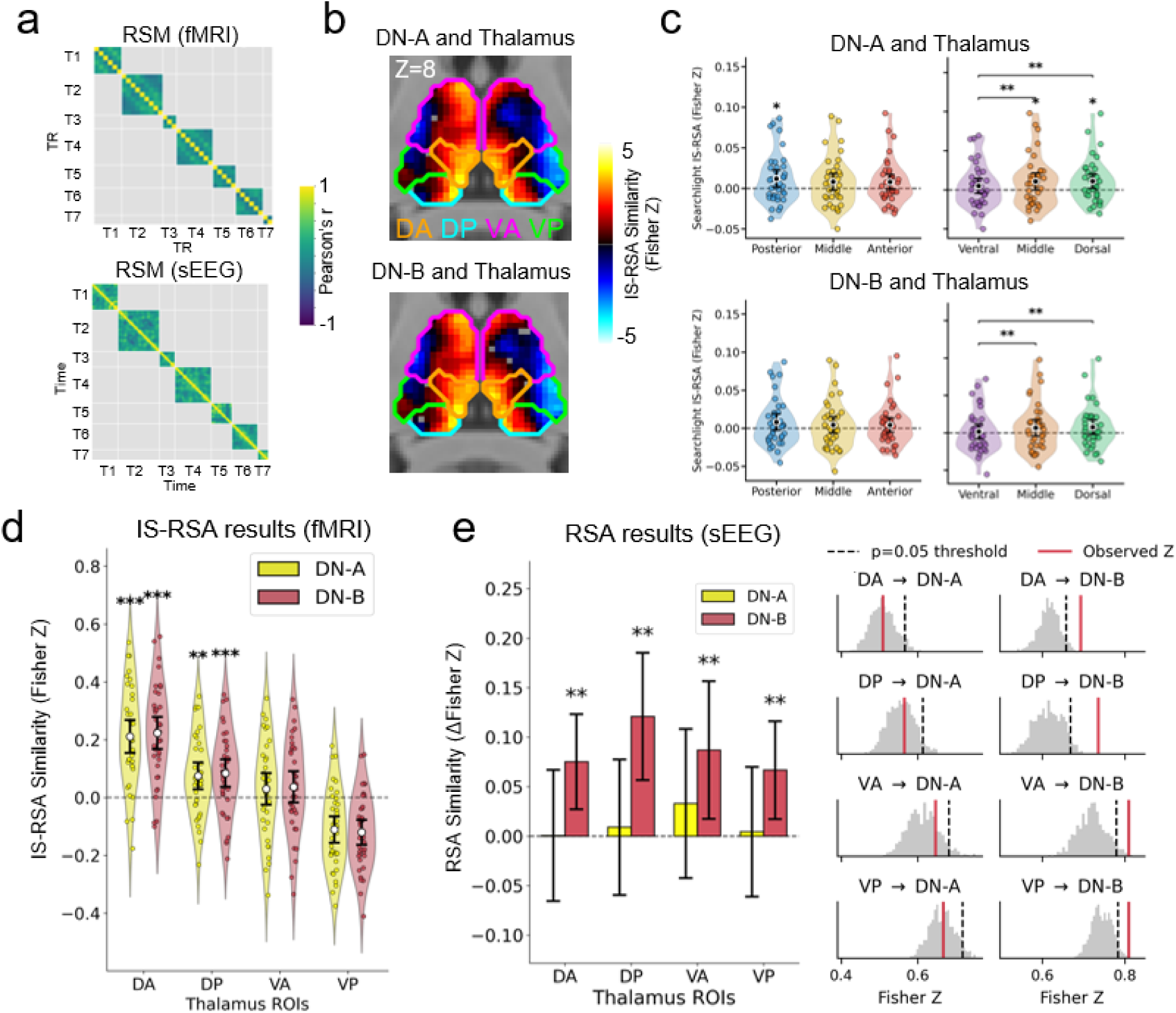
Thalamus shares ToM-related representations with the DN. **(a)** TR-by-TR representational similarity matrices (RSMs) for fMRI (top) and time-by-time RSMs for sEEG (bottom) during ToM events. Each cell in the matrix represents Pearson’s correlation calculated between the spatial activity patterns (across voxels for fMRI or recording sites for sEEG) of two corresponding TRs or time points. **(b)** Fine-grained spatial topography of thalamocortical representational coupling revealed by searchlight inter-subject RSA (IS-RSA) using a leave-one-out (LOO) framework. **(c)** Spatial binning analysis of the searchlight IS-RSA along the anterior-posterior and dorsal-ventral axes. **(d)** ROI-level fMRI IS-RSA coupling results. **(e)** sEEG RSA coupling results. Bar plots (left) show the representational coupling effect, defined as the difference between the true Fisher’s Z and the mean of the null distribution (Δ Fisher’s Z). Histograms (right) illustrate the observed Fisher’s Z values (red solid lines) relative to the permutation-based null distributions (grey) generated via within-event circular shifts. The black dashed lines represent the p = 0.05 significance threshold based on the null distribution. For all the plots, each dot represents an individual subject, and error bars represent 95% confidence intervals. Pairwise comparisons among the three spatial bins in (c) were evaluated using two-tailed paired t-tests, while single-sample tests against zero (including the regional bin-specific tests and all ROI-level inferences) followed a directional one-tailed approach. * *p* < 0.05, ** *p* < 0.01, *** *p* < 0.001 (FDR-corrected).

Next, we conducted a ROI-based analysis using predefined anatomical boundaries to formally evaluate individual subregions (**Fig. 2d**). Consistently, we observed significant positive IS-RSA coupling between dorsal thalamic subregions (both DA and DP) and both DN subsystems across participants (FDR-corrected *p* < 0.05).

We next asked whether this representational alignment generalized to direct electrophysiological signals. To this end, we applied RSA to the sEEG HFA signals, constructing time-by-time RSMs across all ToM events, which revealed block-diagonal structure corresponding to event-specific representations (**Fig. 2a**, bottom). We then quantified representational coupling between thalamic subregions and DN subsystems by correlating their RSMs. All thalamic subregions exhibited significant positive representational coupling with the DN, particularly with DN-B (**Fig. 2e**; FDR-corrected *p* < 0.05). These effects exceeded permutation-based null distributions generated via within-event circular shifts, suggesting that the observed coupling reflects shared ToM-related information rather than spurious autocorrelation.

To explore the underlying causes of the discrepancies observed between fMRI and sEEG results, we conducted further control and exploratory analyses. First, we investigated whether the modality-specific differences in representational coupling were driven by the sparse spatial sampling inherent to intracranial recordings. We performed a control IS-RSA utilizing “virtual electrodes” extracted from the 7T fMRI data based on actual sEEG coordinates (see Methods). This sparse-sampling fMRI analysis yielded results highly consistent with those obtained using the complete cortical networks, preserving the specific representational alignment between the dorsal thalamus and the DN (**Fig. S2**). This confirms that the distinct coupling profiles observed across modalities are not an artifact of spatial sparsity in sEEG recordings. Next, we asked whether these nuances might stem from the intrinsic physiological differences between hemodynamic and electrophysiological signals. To investigate this, we conducted a direct cross-modal temporal correlation analysis between the hemodynamic response function (HRF)-convolved sEEG HFA and the native-space fMRI BOLD signals at corresponding coordinates (see Methods). At the individual electrode level, we observed substantial spatial heterogeneity in this temporal coupling (**Fig. S3A**). At the network level, different regions exhibited markedly divergent neurovascular coupling profiles. Within the DN, only the dorsomedial subsystem (DN-B) demonstrated a significant positive correlation between HFA and BOLD signals at the group level (**Fig. S3B**), thereby partially explaining why our sEEG RSA results predominantly highlighted DN-B. While coupling within DN-A was numerically positive, it did not reach statistical significance. In contrast, the relationship between HFA and BOLD signals within the thalamus was numerically negative across subregions, reaching statistical significance specifically within the VP thalamus.

In sum, despite fMRI and sEEG capturing neural dynamics from different temporal and physiological scales, a consistent cross-modal overlap emerged showing that the thalamus shared ToM-related representation with the DN. In particular, dorsal thalamic regions (DA and DP) exhibited robust representational coupling with DN-B in both fMRI and sEEG, with the cross-modal nuances partially accounted for by region-specific variations in neurovascular coupling.

### Enhanced bidirectional communication in the thalamocortical system during ToM

Having found that the thalamus shares ToM-related representation with the DN, we next sought to examine whether this shared information is accompanied by direct, ToM-modulated interregional communication. To elucidate the directional dynamics of this network, we performed psychophysiological interaction (PPI) analysis, isolating event-dependent changes in effective connectivity during mentalizing across both hemodynamic and electrophysiological modalities.

#### 7T fMRI reveals the spatially comprehensive topography of thalamocortical communication during ToM

To obtain a spatially comprehensive view of the thalamocortical connectivity dynamics during ToM, we first leveraged the whole-brain coverage and high spatial resolution of 7T fMRI using a LOO inter-subject PPI (IS-PPI) framework. By evaluating the task-dependent coupling between a LOO group-averaged physiological state of a seed region and an individual’s target region, this approach allowed us to examine task-modulated communication both at the level of predefined ROIs and across the fine-grained voxel-wise topography within the thalamocortical system.

To map the organization of these interactions on a fine-grained scale, we first conducted voxel-wise IS-PPI mapping within a ROI-masked search space **(Fig. 3a)**. This analysis showed spatial heterogeneity in how different thalamic subregions modulate the DN during mentalizing. Both the anterior and dorsal thalamus (VA, DA and DP) exhibited positive task-modulated PPI across the DN, although the dorsal regions additionally showed a complex, interleaved pattern with some localized negative modulations. In contrast, the VP thalamus exhibited primarily negative task-modulated connectivity with the DN.

**Figure 3.**
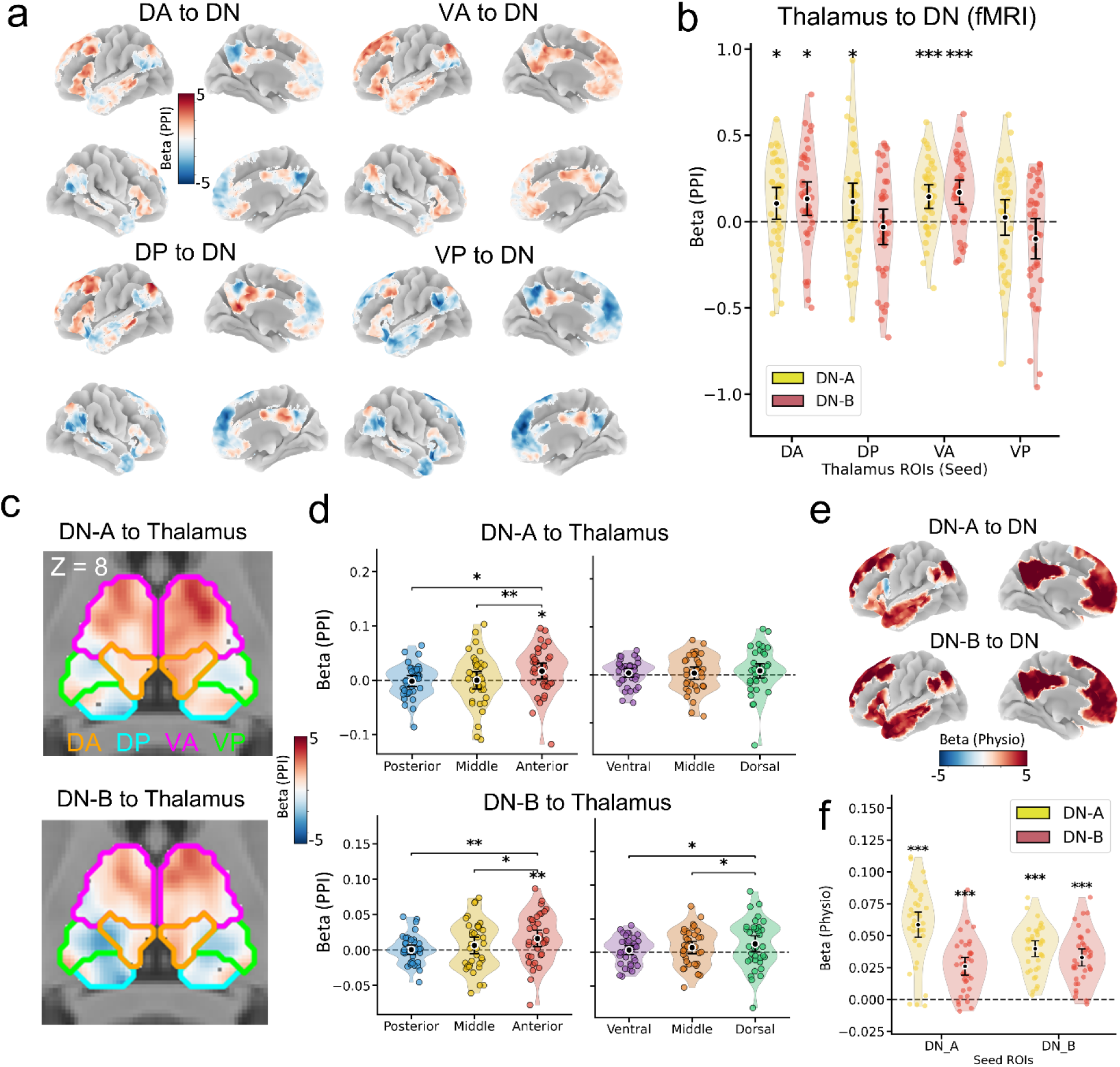
Enhanced bidirectional communication in the human thalamocortical system during ToM using 7T fMRI IS-PPI analysis. Throughout these panels, Beta (PPI) represents ToM-modulated connectivity changes, whereas Beta (Physio) represents intrinsic physiological baseline connectivity. **(a)** Voxel-wise IS-PPI mapping of ToM-modulated thalamic connectivity across the DN. **(b)** ROI-level IS-PPI results for ToM-modulated thalamus-to-DN connectivity. **(c)** Voxel-wise mapping of ToM-modulated corticothalamic (DN-to-thalamus) communications. **(d)** Spatial binning analysis of ToM-modulated corticothalamic modulation along the anterior-posterior (A-P) and dorsal-ventral (D-V) axes. **(e)** Voxel-wise visualization of intrinsic connectivity between DN-A and DN-B. **(f)** ROI-level intrinsic baseline connectivity within DN subsystems. In (b), (d), (f), each dot represents an individual subject. Error bars represent 95% confidence intervals. Pairwise comparisons among the three spatial bins in (d) were evaluated using two-tailed paired t-tests, while single-sample tests against zero (including the regional bin-specific tests and all ROI-level inferences) followed a directional one-tailed approach. * *p* < 0.05, ** *p* < 0.01, *** *p* < 0.001 (FDR-corrected).

Next, we formally quantified task-modulated connectivity at the pre-defined ROI level **(Fig. 3b)**. Consistent with the voxel-wise topography, these analyses revealed that both the anterior and dorsal thalamic subregions exhibited significantly enhanced task-modulated connectivity to the DN. Specifically, the DA and VA subregions showed robust task-dependent coupling with both the DN-A and DN-B subsystems, while the DP exhibited significant coupling selectively with the DN-A.

In the reciprocal corticothalamic (DN-to-thalamus) direction, voxel-wise mapping **(Fig. 3c)** and subsequent spatial binning analysis **(Fig. 3d)** showed highly organized topographic gradients. For both DN subsystems, task-modulated communication to the thalamus followed a significant anterior-posterior (A-P) gradient, where modulation was significantly stronger in the anterior compared to middle and posterior sections **(Fig. 3d)**. Furthermore, along the dorsal-ventral (D-V) axis, the DN-B subsystem exhibited a distinct topographic preference, with task-modulated inputs significantly favoring the dorsal thalamus over the middle and ventral segments **(Fig. 3d)**. This high-resolution hemodynamic evidence underscores that during social inference, the DN precisely targets its feedback to the anterior and dorsal portions of the thalamus. However, task-modulated connectivity at the pre-defined ROI level did not yield statistically significant results **(Fig. S4)**, indicating that functional heterogeneity within these predefined subregions may dilute topographically specific feedback during ROI-level signal extraction.

To understand how thalamic signals are integrated across the whole DN, we hypothesized that the DN’s strong internal coherence facilitates the rapid propagation of localized thalamic inputs. To examine this, we analyzed the intrinsic physiological baseline connectivity of the DN. Analysis of the physiological main effects showed a robust, task-independent connectivity within the DN subsystems **(Fig. 3e, 3f)**, highlighting dense communication between DN-A and DN-B that facilitates the integration of information across the network.

#### Electrophysiological evidence for *enhanced* bidirectional signaling

To validate these hemodynamic observations with direct electrophysiological evidence, we performed PPI analysis of sEEG HFA. During ToM events, both the DA and DP thalamic subregions exhibited significantly increased connectivity to both DN subsystems (FDR-corrected *p* < 0.05) **(Fig. 4a)**. Notably, while this task-dependent modulation was statistically robust within the dorsal regions after multiple-comparison correction, the VA thalamus also exhibited a clear trend toward enhanced connectivity, with 95% confidence intervals excluding zero (95% CI [0.004,0.055] for VA-to-DN-A PPI; 95% CI [0.011,0.049] for VA-to-DN-B PPI). In contrast, the VP subregion showed no such trend, with its intervals spanning zero (95% CI [-0.032, 0.030] for VP-to-DN-A PPI; 95% CI [-0.008,0.041] for VP-to-DN-B PPI). This electrophysiological signaling pattern selectively mirrors the higher-order thalamic engagement observed in our 7T fMRI dataset.

**Figure 4.**
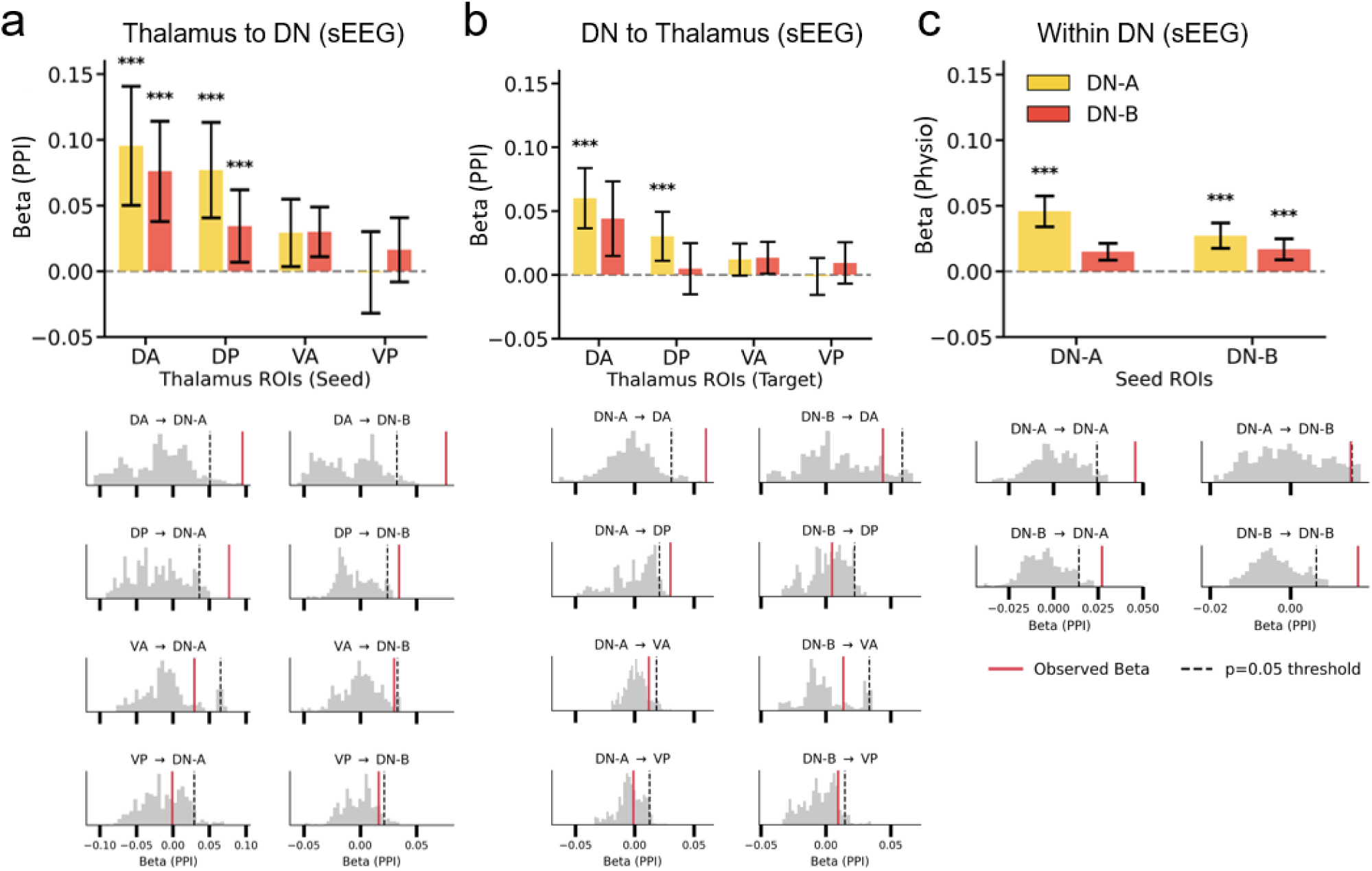
Enhanced bidirectional communication in the human thalamocortical system during ToM using sEEG PPI analysis. Throughout these panels, Beta (PPI) represents ToM-modulated connectivity changes, whereas Beta (Physio) represents intrinsic physiological baseline connectivity. **(a)** ToM-modulated connectivity from the thalamus to the DN. **(b)** Reciprocal ToM-modulated connectivity from the DN to the thalamus. **(c)** Intrinsic physiological connectivity within the DN subsystems. Error bars represent 95% confidence intervals. Beneath each bar plot, the corresponding permutation-based null distributions are displayed as grey histograms. The red solid lines indicate the observed Beta values, and the black dashed lines denote the one-tailed *p* = 0.05 significance threshold. All ROI-level inferences followed a directional one-tailed approach. *** *p* < 0.001 (FDR-corrected).

Next, our examination of corticothalamic (DN-to-thalamus) pathways showed a complementary return loop. Aligning with the dorsal-specific feedback observed in our fMRI data, ToM-modulated connectivity directed back to the dorsal thalamus was predominantly driven by the DN-A, which demonstrated significantly enhanced effective connectivity to both the DA and DP subregions (FDR-corrected *p* < 0.05) **(Fig. 4b)**. While directed communication from the DN-B did not exhibit significant ToM-related modulation to any thalamic subregion after multiple-comparison correction, we observed a clear trend of enhanced connectivity selectively targeting the DA and VA subregions, with their 95% CI excluding zero (95% CI [0.015, 0.073] for DN-B to DA; 95% CI [0.001, 0.026] for DN-B to VA). This electrophysiological trend closely mirrors the topographically specific corticothalamic feedback from the DN-B to the anterior thalamus observed in our 7T fMRI data, supporting convergent cross-modal architecture. Consistent with the fMRI baseline results, analysis of the physiological main effects in sEEG confirmed strong baseline intrinsic connectivity binding the cortical DN subsystems **(Fig. 4c)**. Specifically, while the baseline connectivity from DN-B to DN-A was robustly significant, the reciprocal pathway from DN-A to DN-B exhibited a slightly weaker but clear trend, with its uncorrected 95% CI excluding zero (95% CI [0.009, 0.021]). This indicates persistent, albeit somewhat asymmetric, information exchange within the DN. Collectively, these sEEG findings support a specialized bidirectional circuit where the dorsal and anterior thalamus broadcasts ToM-related signals across the DN while receiving feedback mainly from its core subsystem.

These convergent findings across hemodynamic and electrophysiological scales reveal that despite lacking canonical local activation during mentalizing, the thalamus, particularly its dorsal and anterior subregions, exhibits enhanced, task-modulated bidirectional connectivity with the DN. This bidirectional architecture, characterized by anatomically specific thalamocortical signaling and topographically organized corticothalamic feedback, demonstrates the pathways through which the thalamus interacts with cortical networks during social computation.

### Thalamic low-frequency phase unidirectionally gates DN HFA and representational quality

To investigate the neurophysiological mechanisms underlying this directed thalamocortical communication, we examined cross-frequency PAC, which serves as a canonical process for distributed neural coordination wherein low-frequency oscillations from a regulatory hub temporally structure the HFA of a target region. By computing PAC across a comprehensive frequency-frequency space during ToM events relative to implicit baseline, we found broad, significant increases in PAC between the thalamus low-frequency phase (∼ 3-10Hz) and the DN HFA (>70 Hz) **(Fig. 5a)**, with the sole exemption of the VP to DN-A pair. In contrast, an examination of the reverse direction (e.g., DN low-frequency phase modulating thalamic HFA) yielded no significant coupling across any ROI pairs **(Fig. 5b)**. We also examined local PAC within thalamic or DN ROIs (e.g., DN low-frequency phase modulating DN HFA) but found no significant coupling enhancements **(Fig. S5)**.

**Figure 5.**
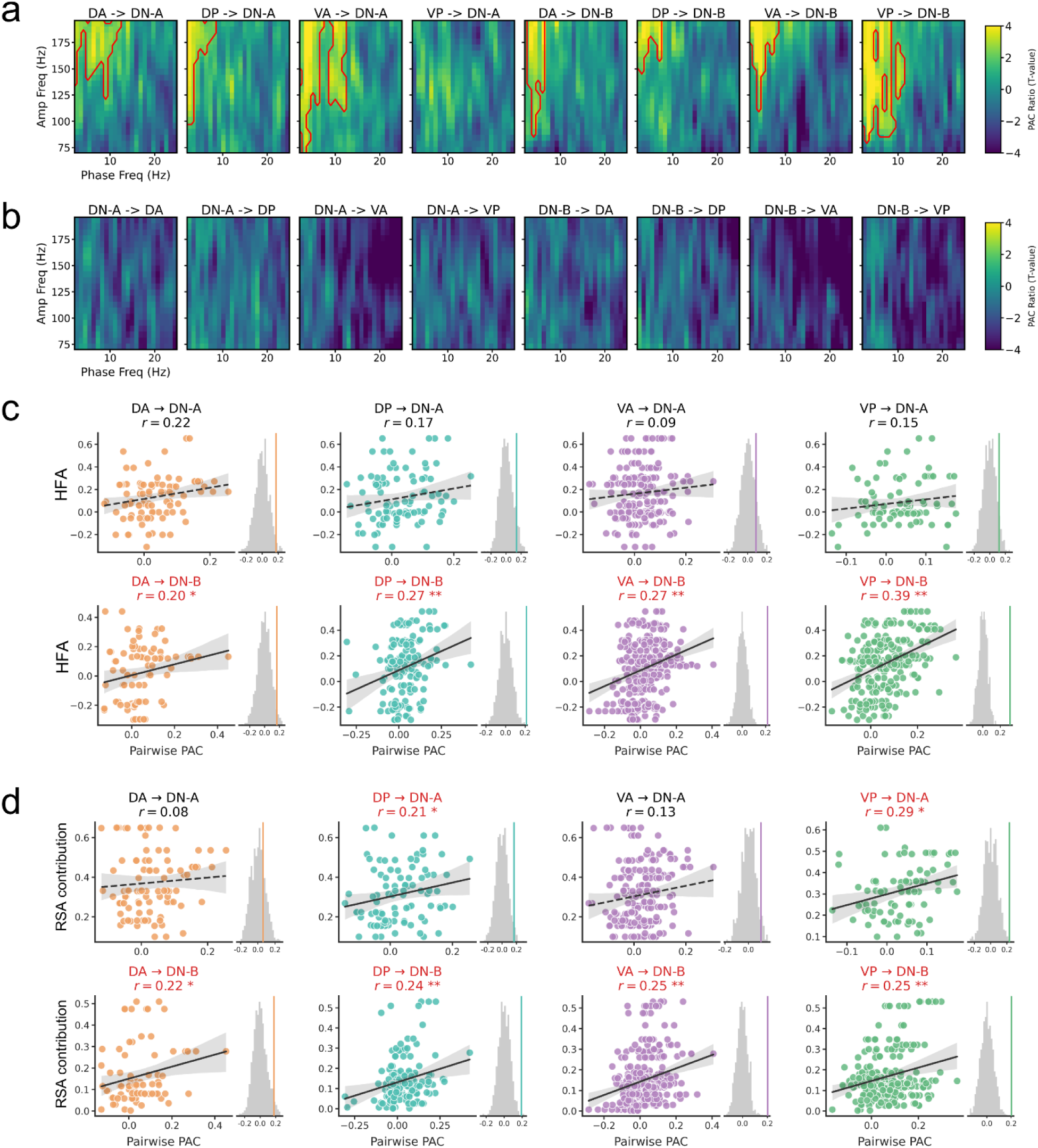
Thalamic low-frequency phase unidirectionally gates DN activation and representation quality. **(a)** ToM-related enhancement of cross-frequency phase-amplitude coupling (PAC) from four thalamic subregions to default network (DN) subsystems (DN-A and DN-B). Color maps represent the group-level normalized PAC contrast ratio (comparing ToM events against non-ToM baseline periods, expressed as T-values) across low-frequency phase (2–25 Hz) and high-frequency amplitude (70–200 Hz). Red outlines denote significant frequency-frequency clusters representing PAC enhancements during mentalizing (one-sample cluster-based permutation test against zero, *p* < 0.05). **(b)** Reciprocal PAC analysis from DN subsystems to thalamic subregions, showing a complete lack of significant ToM-related coupling enhancements in the corticothalamic direction. **(c)** PAC-HFA correlation. Pearson correlations between pairwise thalamic PAC (coupling between 3–10 Hz thalamic phase and 70–170 Hz DN amplitude) and the local HFA of target cortical sites during ToM. Each point represents an individual thalamocortical electrode pair. Significant positive correlations (highlighted in red with solid trendlines) demonstrate that thalamic modulatory inputs strongly scale local cortical excitability. Adjacent to each scatter plot is the null distribution of correlation coefficients generated via 1000 global permutations (gray histogram). The colored solid vertical line indicates the observed correlation coefficient. **(d)** PAC-representation correlation. Pearson correlations between pairwise thalamic PAC and RSA contribution scores of the target cortical sites. The RSA contribution quantifies the impact of a “virtual lesion” of the target site on the network-level representational geometry. Null distributions from the permutation tests are displayed to the right of each scatter plot, identical to (c). * *p* < 0.05, ** *p* < 0.01, *** *p* < 0.001, after FDR correction.

Furthermore, we asked whether this unidirectional thalamic pacing drives the functional execution of local cortical processing. To quantify the modulatory drive received by the DN, we evaluated the pairwise PAC for every valid thalamocortical connection. We then correlated this incoming modulatory input with the target (DN) site’s local task-evoked activation (HFA). The strength of pairwise thalamocortical PAC significantly and positively predicted the HFA levels of individual DN sites. Notably, this gating effect was present across the DN-B, where PAC inputs from all four thalamic subregions significantly scaled local cortical activation (with *r* values ranging from 0.20 to 0.39, permutation-based FDR-corrected *p*s < 0.05) **(Fig. 5c)**.

We next determined if thalamic regulation extends beyond modulating activation levels to shape the multivariate representation processed by the cortex. To quantify the representational quality of each DN site, we implemented a virtual lesion approach based on RSA to calculate an RSA contribution score for individual sites (see Methods). Specifically, we measured the extent to which the network’s overall ToM-related representational geometry degraded when a given site was systematically excluded **(Fig. 5e)**. We found that the strength of pairwise thalamic PAC significantly predicted a cortical target site’s RSA contribution score. While the DP and VP subregions showed significant modulatory effects on the DN-A subsystem (*r* = 0.21 and 0.29, respectively), this relationship was once again observed across the targeted DN-B subsystem, where PAC from all four thalamic subregions significantly predicted the RSA contribution scores of individual sites (with r values ranging from 0.22 to 0.25, permutation-based FDR-corrected *p*s < 0.05) **(Fig. 5d)**.

Together, these results provide direct physiological evidence for the human thalamus as an orchestrator during mentalizing. Although the thalamus does not exhibit overt local task-evoked activation itself, it utilizes unidirectional low-frequency phase dynamics to rhythmically gate both the localized activation strength and the representational quality of the DN to support mentalization.

## Discussion

For decades, conventional models of ToM have primarily operated within a corticocentric framework that positions the DN as the central core of mentalizing ^2,10–13^. While our findings confirm that the DN remains the primary cortical substrate for the foreground execution of naturalistic mentalizing, they demonstrate that the execution of these high-level cortical processes is integrally tied to subcortical orchestration. Integrating high-field 7T fMRI and direct intracranial sEEG recordings, we demonstrate a critical role of thalamic-DN interactions in naturalistic ToM. Specifically, we observed a clear functional dissociation within the thalamocortical loop: while the DN underpins the foreground activation during ToM, the locally “silent” thalamus acts as a background regulator that participates in mentalizing through shared representational geometry and heightened bidirectional connectivity with DN hubs. Furthermore, this interregional coordination is sustained by a unidirectional, asymmetric rhythmic regulation process, whereby the low-frequency phase of the thalamus gates both the local activation magnitude and the multivariate representational quality within the DN, particularly its dorsomedial DN-B subsystem. This circuit-level integration supports a shift in how we characterize the neural architecture of social behavior: a predominantly DN-localized model of mentalizing toward a thalamocortical framework. Extending prior evidence from non-human animal models and non-social cognitive domains, our findings establish, through convergent human multimodal evidence, the thalamus as an important coordinator of social cognition.

This coordinated interaction demonstrates that the human thalamus can exert a profound regulatory influence on high-level social networks without relying on canonical, task-related local activation. In univariate neuroimaging experiments, the absence of overt univariate task activation may lead to the assumption of functional irrelevance, often peripheralizing subcortical structures in empirical characterizations of high-level cognition. However, increasing insights into higher-order thalamic nuclei involvement in cognition challenge this cortico-centric view, positioning these structures as flexible coordinators that shape cortical dynamics without necessarily inducing substantial changes in their own metabolic or local firing-rate baselines ^17,19,24,32^. Prior work in non-human animal models and computational modeling demonstrate that higher-order thalamic hubs, such as the mediodorsal or anterior thalamus, sustain cortical behavioral representations not by driving localized activation, but by amplifying intrinsic cortical connectivity and modulating long-range oscillatory coherence ^28,29,32,33,48^. Our findings extend this regulatory principle into the domain of complex social cognition in humans, indicating that the human thalamus participates in mentalizing through an alternative computational mode centered on distributed neural coordination and network-level synchronization.

Anatomically, the thalamic-DN coordination observed during naturalistic ToM is rooted in the intrinsic and extensive anatomical and functional connectivity pathways linking the thalamus to the neocortex ^19,26,49–52^. Crucially, our findings demonstrate that these thalamic-DN pathways are not static relays, but instead undergo dynamic, task-dependent enhancement during naturalistic mentalizing, suggesting that the thalamocortical network flexibly reconfigures to meet the high-level computational demands of social cognition. This network-level modulation exhibits a highly organized functional anatomy within the thalamic subdivisions. Our observation that the dorsal and anterior thalamic subregions exhibit close representational alignment and functional connectivity with the DN during ToM events aligns with structural and functional mapping, which highlights dense, reciprocal wiring between these higher-order nuclei (such as the mediodorsal and anterior groups) and prefrontal association areas, including core nodes of the DN ^21,25,51,53–55^. In contrast, the VP subregion remained largely unengaged during ToM processing. Because the VP thalamus is classically characterized as a first-order sensorimotor relay, it is primarily associated with processing low-level sensory signals rather than supporting abstract cognitive or social inferences ^17,18^. Thus, these subregional variations underscore that the subcortical contribution to social cognition does not reflect a diffuse, generalized thalamic change; instead, it relies on an anatomically specific thalamocortical framework tailored to high-level associative processing.

The neurophysiological basis of this thalamocortical communication is further illuminated by the cross-frequency PAC results. PAC serves as a canonical mechanism for distributed neural coordination, wherein HFA of a target region is temporally organized by the low-frequency phase of a regulatory hub, providing directed rhythmic modulation ^36,56–58^. In this dynamic framework, HFA is widely recognized as a robust proxy for local population spiking and localized cortical computation, whereas low-frequency neural oscillations are uniquely suited to facilitate long-range communication and integrate distributed networks ^59–61^. For example, a recent study ^34^ provided direct intracranial evidence that human higher-order thalamic nuclei gate conscious perception via thalamofrontal PAC, utilizing low-frequency phase to rhythmically modulate prefrontal HFA. Our findings extend this critical subcortical gating mechanism far beyond sensory awareness, positioning it as a critical mechanism supporting abstract social cognition. During naturalistic mentalizing, we observed a robust unidirectional PAC, with the thalamic low-frequency phase pacing the DN HFA, yet no significant coupling in the reverse direction. This distinct asymmetry positions the thalamus as an active rhythmic gatekeeper that temporally coordinates foreground cortical operations. Crucially, this modulation transcends mere epiphenomenal synchrony. We demonstrate that the strength of the incoming thalamocortical PAC directly predicts both the localized activation magnitude and the multivariate representational quality of the cortical target sites, linking subcortical pacing to the fidelity of cortical computation. Notably, this gating effect was most pervasive across the dorsomedial DN-B subsystem, where modulatory inputs from all evaluated thalamic subregions significantly scaled local processing metrics. This pronounced susceptibility implies that DN-B may function as a primary cortical locus for social inference, relying on thalamic temporal coordination to assemble and maintain the complex neural representations required for mentalizing.

Although our multimodal approach suggested a core thalamocortical alignment during naturalistic ToM, subtle discrepancies emerged between the fMRI and sEEG findings. For example, while 7T fMRI revealed broad representational coupling across the dorsal thalamus and both DN subsystems, sEEG representations were predominantly centered on the DN-B. While prior multimodal investigations (including studies using common naturalistic stimuli) have demonstrated a positive correlation between cortical HFA and the BOLD signal ^62,63^, growing evidence suggests that this HFA-BOLD coupling relationship is not uniformly distributed but varies across brain regions ^64,65^. Our supplementary analyses first effectively rule out the spatial sparsity of intracranial sampling as a confounding artifact, and then demonstrated that these cross-modal nuances may indeed stem from intrinsic physiological differences between hemodynamic and electrophysiological signals for different regions. Within the cortical DN, a robust, significant positive HFA-BOLD temporal correlation was selectively observed in the DN-B, providing a potential physiological explanation for the heightened prominence of DN-B in our sEEG RSA results. In stark contrast, the temporal relationship within the thalamus trended numerically negatively across subregions, reaching statistical significance in the VP subregion. These divergent coupling profiles indicate that fMRI and sEEG HFA may capture partially distinct yet complementary windows into thalamocortical dynamics. By integrating high-field 7T fMRI and direct intracranial recordings, our dual-modality framework leverages these complementary axes to offer a more nuanced and multi-layered characterization of subcortical regulation during human social cognition.

Our results offer practical perspectives for understanding neurodevelopmental and psychiatric conditions characterized by social cognitive difficulties, such as autism spectrum disorder and schizophrenia ^42,66,67^. While traditional models of these conditions have focused primarily on localized cortical impairments, our data support a circuit-level view suggesting that certain symptoms may stem from altered thalamocortical communication ^68,69^. Characterizing these impairments within a thalamocortical framework may aid the development of targeted therapeutic interventions. In this context, advanced non-invasive neuromodulation techniques present promising avenues for targeting deep brain structures without the constraints of surgical implantation. For example, emerging modalities such as transcranial ultrasound stimulation (TUS) ^70,71^ and transcranial temporal interference stimulation (TIS) ^72,73^ have demonstrated the capacity to focus stimulation onto subcortical targets, including specific thalamic nuclei, with high spatial precision. Utilizing TUS or TIS to non-invasively modulate the thalamic low-frequency rhythms identified here might offer a practical path toward modulating downstream cortical computation and representational quality within the cortical network.

## Conclusion

We describe a thalamic-DN circuit architecture supporting human social cognition. By demonstrating that the thalamus rhythmically gates the DN during mentalizing without displaying localized, task-related activation, we highlight its role as a background regulatory hub of foreground cortical computations. These observations suggest that models of higher-order social cognition could be expanded to incorporate circuit-level thalamocortical dynamics, moving beyond an exclusively cortical focus to better understand how distributed brain networks organize human social thought.

## Methods

### Participants

Thirty-seven healthy adults (24 females, 13 males; age range: 18-35y, mean age: 24.08, standard deviation: 3.98) participated in the 7T fMRI study. All participants were right-handed, with normal or corrected-to-normal vision, and no reported history of neurological or psychiatric conditions. All scanning was conducted at the McConnell Brain Imaging Centre at the Montreal Neurological Institute. Prior to participation, all individuals were screened for 7T MRI contraindications and provided written informed consent consistent with the study protocol approved by the McGill University Institutional Review Board. All procedures complied with the ethical standards of the 1964 Declaration of Helsinki.

Eight presurgical epilepsy patients (5 females, 3 males; age range: 22-67, mean age: 39.88, standard deviation: 15.36) participated in this sEEG study. Electrode implantation and placement were determined solely by clinical considerations for localization of the seizure onset zone. The study protocol was approved by the Stanford University Institutional Review Board, and all participants provided written informed consent before taking part in the research procedures.

### Experimental stimuli

Participants in both the 7T fMRI and sEEG cohorts viewed “Partly Cloudy” ^41^, a 5.6-minute animated short film. The movie depicts the evolving friendship between a stork and a cloud through a series of social interactions, and it has served as a well-validated paradigm for studying spontaneous mentalizing ^8,9,42,74^. For our analysis, we utilized the initial 313-second segment of the movie from its onset, excluding only the end credits. To identify ToM-related neural processing, we adopted the temporal annotations established by Richardson et al. ^8^. These annotations define seven distinct ToM events (T1–T7) totaling 68 seconds of the film, with individual event durations ranging from 4 to 16 seconds. Detailed timing and descriptions for each event are provided in Supplementary Table 1.

### Parcellation and definition of ROIs

To delineate cortical and subcortical regions of interest (ROIs), we applied the Schaefer 400-region parcellation ^43^ with Yeo 17-network framework ^44^ for cortical parcellation. We specifically focused on two subsystems of the DN that are engaged during mentalizing: the DN-A (core subsystem) and the DN-B (dorsal medial subsystem). For the human thalamus, we employed the Melbourne Subcortex Atlas (Scale 2) ^45^, which parcellates the thalamus into four ROIs: dorsal anterior (DA), dorsal posterior (DP), ventral anterior (VA), and ventral posterior (VP). We applied these parcellation schemes and ROI definitions consistently to analyze both the sEEG and 7T fMRI datasets

### 7T multi-echo fMRI acquisition and preprocessing

7T anatomical and functional neuroimaging data were acquired with a Siemens Magnetom Terra 7T scanner at the McConnell Brain Imaging Centre at the Montreal Neurological Institute. Anatomical images were acquired using a T1-weighted MPRAGE (0.8 mm isovoxels; TR = 3300ms; TE = 2.74ms; FOV = 230mm; 224 sagittal slices; TI = 1100ms; flip angle = 4°; echo spacing = 7.4ms). Multi-echo fMRI data were acquired with a 2D BOLD echo-planar imaging sequence (1.9 mm isotropic voxels, TR = 1.72 s, TE’s = 11.2 / 27.8 / 44.4 ms, flip angle = 67°, FOV = 224mm, 75 slices, multiband factor = 3, iPAT GRAPPA = 3; Phase partial Fourier 6/8, echo spacing = 0.53 ms).

Multi-echo fMRI data were preprocessed and denoised using the multi-echo independent component analysis (ME-ICA) pipeline v3.9.3 ^75,76^. ME-ICA acts as an integrated workflow that leverages the TE-dependence model of the BOLD signal to optimally combine multi-echo data and robustly separate neural activity from non-BOLD sources of noise. All task fMRI analyses were conducted using the multi-echo denoised and anatomically co-registered voxel-wise time series (T1C_MEDN outputs).

### fMRI analyses

#### BOLD activation analysis

We performed first-level GLM analysis for each of the 37 participants using the Nilearn library ^77^. For each participant, we truncated the functional time series to the duration of the movie (313 s) and modeled the ToM events as block regressors based on seven temporally precise events (totaling ∼68s) from the “Partly Cloudy” film, as annotated in previous studies ^8^, with the remaining periods of the movie serving as an implicit baseline to control for continuous sensory processing. We convolved these event timings with a canonical double-gamma hemodynamic response function (HRF). To minimize low-frequency drifts, we included a cosine drift model with a high-pass filter cutoff of 0.01 Hz. We also accounted for temporal autocorrelation using a first-order autoregressive (AR(1)) noise model and standardized the voxel time series before fitting the model.

We implemented a robust registration pipeline using Advanced Normalization Tools (ANTs) ^78^ to transform native functional maps into MNI152 space. First, we performed N4 bias field correction on the T1-weighted anatomical images. We then computed a rigid-body transformation to align the mean functional (BOLD) image to the individual T1-weighted image. Subsequently, we utilized the Symmetric Normalization (SyN) algorithm to estimate the non-linear warp from the participant’s anatomical space to the MNI152 template. We concatenated these transformations to resample the native Z-score and beta maps into 2mm isotropic MNI space.

We performed group-level analysis using a second-level one-sample t-test on the normalized beta maps. We applied a 2mm FWHM Gaussian smoothing kernel to the group-level estimates to enhance the signal-to-noise ratio. To identify significant activations, we employed false discovery rate (FDR) correction (*q* < 0.05, one-tailed *Z* > 0) with a cluster-extent threshold of 10 voxels. For targeted investigations of the thalamus, FDR correction was performed within the whole thalamus (with a cluster-extent threshold of 5 voxels). Finally, we conducted ROI analysis by extracting mean Z-scores from these ROIs and performed one-sample t-tests (FDR-corrected) to verify network-level engagement during mentalizing.

#### fMRI IS-RSA analysis

We characterized the sustained representational shared information between the thalamus and the DN using a leave-one-out (LOO) inter-subject RSA (IS-RSA) approach. For each ROI, we first Z-scored the voxel-wise BOLD time series across both temporal and spatial axes to remove static anatomical baselines and global signal fluctuations. We constructed TR-by-TR RSMs for each ToM event by calculating the Pearson correlation across voxels for every pair of TRs, accounting for a canonical HRF delay of 3 TRs (∼5.16s). To reliably quantify the coupling between ROIs, we employed a bidirectional symmetric approach: for each participant, we matched their individual thalamic RSM with the group-averaged DN RSM of all other participants (Direction 1), and vice versa (Direction 2). The upper triangular vectors of these RSMs were concatenated across all valid ToM events, and the coupling strength was calculated as the Pearson correlation between individual and group-averaged vectors. After applying a Fisher’s Z transformation, the two directional coupling scores were averaged for each subject to provide a robust measure of symmetric representational sharing. To assess significance, we performed a one-tailed one-sample t-test comparing the group-level Fisher’s Z against 0 and applied FDR correction across all thalamocortical pathways (q < 0.05).

To map the fine-grained spatial topography of these shared representations within the thalamus, we extended our IS-RSA framework using a volumetric searchlight approach. For each participant, analyses were conducted in their native anatomical space to preserve spatial precision. We defined a spherical searchlight with a 4 mm radius centered on every voxel within the native-space whole-thalamus mask. Voxel-wise BOLD time series within each sphere were normalized (Z-scored) and concatenated across all ToM events, accounting for a canonical HRF delay of 3 TRs. For each local thalamic sphere, we computed a TR-by-TR RSM using Pearson correlation across voxels. Consistent with the ROI-level analysis, we applied a LOO framework to isolate stimulus-locked representational structures. The representational coupling for each searchlight sphere was quantified by calculating the Pearson correlation between the lower triangular vector of the sphere’s RSM and the group-averaged TR-by-TR RSM of the target cortical subsystem (DN-A or DN-B) derived from all other participants. The resulting native-space correlation maps were Fisher Z-transformed. These statistical maps were subsequently normalized to the MNI152 space using ANTs. Finally, we performed a second-level one-sample t-test on the normalized maps.

To quantitatively evaluate spatial gradients in thalamocortical representational coupling, we applied a volume-equated spatial binning analysis to the normalized searchlight IS-RSA maps. The thalamic mask was parcellated into three equal-volume terciles along both the anterior-posterior (Y-axis) and dorsal-ventral (Z-axis) spatial axes, using the 33.3rd and 66.7th percentiles of their respective MNI coordinates. For each subject, the mean Fisher Z-transformed coupling value was extracted from each spatial bin. Group-level topological gradients were assessed by comparing the extracted values across bins using repeated-measures paired t-tests, while regional significance against zero was evaluated using one-tailed one-sample t-tests, with all *p*-values FDR-corrected (*q* < 0.05).

To ensure that subsequent cross-modal comparisons between fMRI and SEEG modalities are not merely driven by the sparse spatial sampling inherent to intracranial recordings, we conducted a sparse-sampling control IS-RSA. Specifically, we constructed “virtual electrode” atlases for the DN-A and DN-B subsystems by generating 4-mm radius spherical ROIs centered on the exact MNI coordinates of all valid sEEG recording sites. These sparse MNI atlases were then mapped back to each participant’s native functional space using the inverse of the non-linear SyN warps and affine transformations generated via ANTs. Voxel-wise BOLD time series were extracted exclusively from these subject-specific sparse cortical masks. The subsequent construction of TR-by-TR RSMs and the LOO IS-RSA between these sparse cortical subsystems and the thalamic subregions were performed using the exact same procedures as described above.

#### fMRI IS-PPI

We evaluated task-modulated directional connectivity using a hybrid inter-subject PPI (IS-PPI) framework. Traditional within-subject PPI is often confounded by shared, non-neural physiological noise (e.g., respiration, cardiac rhythms) and intrinsic autocorrelations. To overcome this limitation and isolate stimulus-locked, task-specific neural communication, we implemented an IS-PPI approach utilizing a LOO group-average strategy. We applied this framework across two complementary spatial scales to elucidate the interactions between the thalamus and the DN (or within the DN): a ROI level analysis for evaluating the specific directional coupling between predefined sub-networks, and a voxel-wise analysis to map the fine-grained spatial topography of these interactions.

For the ROI-wise IS-PPI analysis, we first applied a Principal Component Analysis (PCA) across all voxels within the native mask of each region and extracted the first principal component (PC1) to obtain robust representative time series for each ROI. We ensured the sign of the PC1 was consistent with the mean signal of the ROI. For each subject, all analyses were performed within their native anatomical space to preserve spatial precision and avoid interpolation artifacts introduced by standard space warping. To test the hypothesis of directional modulation (e.g., thalamus to DN), we defined the target region (e.g., DN-A) using the individual subject’s actual PC1 time series (Y). The seed region (physiological variable) was defined by computing the group-average PC1 time series from all *other* subjects (the LOO strategy), thus reflecting the shared, stimulus-driven physiological state of the seed independent of the target subject’s idiosyncratic noise. The psychological variable was constructed based on the binary temporal boundaries of the ToM events, convolved with a canonical HRF and subsequently mean-centered. The interaction term (PPI) was generated by multiplying the mean-centered psychological variable with the standardized (Z-scored) LOO physiological time series. We then constructed a GLM for each subject to predict the target Y. The resulting beta weights for the PPI term represented the task-modulated connectivity, and the resulting beta weights for the psychological term represented the intrinsic baseline connectivity. Group-level significance was assessed using one-tailed one-sample t-tests against zero, followed by FDR correction for multiple comparisons across all ROI pairs (*q* < 0.05).

For the voxel-wise IS-PPI analysis, the seed (physiological variable) was extracted as the 1D PC1 time series of the relevant ROI averaged across all other subjects (LOO), while the target was the subject’s 4D native functional time series. To optimize computational efficiency and restrict the search space to relevant anatomy, the first-level GLM was masked using the individual’s native target mask (either the combined thalamus mask or the combined DN mask). The design matrix, identical in construction to the ROI-wise analysis (containing the HRF-convolved psychological variable, the standardized LOO seed physiological variable, and their interaction PPI), was fitted to every voxel within the target mask. This yielded a 3D beta map representing the voxel-wise PPI interaction strength in the subject’s native space. To facilitate group-level visualization, these native 3D beta maps were subsequently warped to the standard MNI template space using ANTs. For group-level inference, a second-level GLM was performed on the MNI-warped beta maps. When targeting the cortex, the resulting group Z-statistic maps were projected onto the *fsaverage* cortical surface to visualize the distribution of connectivity within the DN. Conversely, when targeting the thalamus, group Z-maps were visualized using slices to examine intra-thalamic loci.

Finally, to quantitatively evaluate spatial gradients in DN-to-thalamus modulation, we applied a volume-equated spatial binning analysis. The thalamic mask was parcellated into three equal-volume terciles along both the anterior-posterior (Y-axis) and dorsal-ventral (Z-axis) spatial axes, using the 33.3rd and 66.7th percentiles of their respective MNI coordinates. For each subject, the mean task-modulated IS-PPI beta weight was extracted from each spatial bin. Group-level topological gradients were assessed by comparing the extracted beta weights across adjacent and extreme bins (e.g., posterior vs. middle vs. anterior) using repeated-measures paired t-tests, with *p*-values FDR-corrected (*q* < 0.05).

#### Temporal signal-to-noise ratio (tSNR) analysis

To evaluate the signal quality of the 7T multi-echo fMRI data across our target regions, we computed the voxel-wise tSNR for each participant. The tSNR was calculated in native functional space by dividing the mean of the BOLD time series over the duration of the movie viewing by its temporal standard deviation. To evaluate tSNR across standardized anatomical structures, these native tSNR maps were subsequently warped into the MNI template space using the exact same rigid and non-linear (SyN) transformations generated via ANTs during the GLM analysis. Finally, we generated a group-averaged tSNR map to visualize the overall signal fidelity across the thalamocortical system.

### sEEG recording and electrode anatomical localization

The sEEG signals were recorded using depth electrodes. Seven participants were implanted with AdTech electrodes with contacts spaced 3 or 5 mm apart and a diameter of 0.86 mm, and one participant was implanted with PMT electrodes with contacts spaced 3.5 mm apart and a diameter of 0.8 mm. Intracranial signals were acquired using a Nihon Kohden recording system at a sampling rate of 1000 Hz during movie viewing.

For anatomical localization, a preimplantation T1-weighted structural MRI scan was acquired using a GE SIGNA 3T MRI scanner, and a postoperative computed tomography (CT) scan was obtained following electrode implantation. The postoperative CT image was rigidly co-registered to the preimplantation T1-weighted image, and individual electrode contacts were localized in native anatomical space using BioImage Suite. Cortical surfaces were reconstructed from each participant’s T1-weighted image using FreeSurfer. Electrode coordinates were subsequently transformed from native anatomical space to standard MNI space. Cortical recording sites were assigned to functional networks by mapping their standardized coordinates to the corresponding atlas labels in the Schaefer 400-region parcellation with Yeo 17-network framework ^43^, whereas thalamic sites were assigned to 4 subregions using the Melbourne Subcortex Atlas ^45^.

### sEEG preprocessing

Raw sEEG data were preprocessed using the MNE-Python software package. For each participant, continuous recordings were first temporally synchronized with the movie presentation and cropped to match the precise onset and offset timestamps of the film. Artifactual, epileptic, or excessively noisy channels were identified through clinical logs and visual inspection, and were excluded from downstream processing. The remaining electrode contacts were re-referenced using a local bipolar montage configuration, where the signal from each individual contact was subtracted from its immediate anatomical neighbor along the same electrode shaft. Following re-referencing, a notch filter was applied at 60, 120, and 180 Hz to remove power line noise and its corresponding harmonics.

### sEEG analyses

#### HFA analysis

We extracted high-frequency activity (HFA) from the sEEG recordings as a proxy for local neural processing at each recorded site. We first applied a bandpass filter (70–170 Hz) to the sEEG data and utilized the Hilbert transform to obtain the analytic signal amplitude. We then calculated the instantaneous power by squaring this amplitude, followed by a Gaussian smooth (FWHM = 1 s) across the time dimension. Then, we performed Z-score normalization for each site across the entire duration of the movie.

To safeguard the HFA signal against large transient artifacts and interictal discharges, we implemented an automated detection pipeline based on the median absolute deviation (MAD). We identified any time points exceeding a threshold of 20 MADs as artifacts. To prevent these spike-like artifacts from contaminating the signal during the smoothing process, we masked the detected artifacts along with a ±250 ms buffer. We temporarily filled these masked segments with the median power value of the respective site during the smoothing step, subsequently restoring the mask by setting these periods to NaN for all downstream event-related analyses.

We segmented the continuous HFA time series into seven ToM event windows according to the annotations. For each ROI, we averaged the HFA values across all constituent sites and ToM events to characterize the regional response. At the group level, we evaluated whether specific networks exhibited significant HFA increases during mentalizing using a one-sample sign-flipping permutation test (1000 iterations) implicit movie baseline (zero). This statistical inference followed a one-tailed approach to specifically test for significant task-evoked activations. Finally, we applied FDR correction to account for multiple comparisons across cortical and thalamic networks, setting the significance threshold at q < 0.05.

#### sEEG RSA analysis

We characterized the shared information structure between the thalamus and the DN using RSA based on normalized HFA time series described above. For each ROI, we pooled all available sites across subjects and constructed time-by-time RSMs at a 10 Hz resolution. We generated these RSMs for each ToM event by calculating the Pearson correlation across sites for every pair of time points, resulting in a block-diagonal matrix that captures the internal representational dynamics of mentalizing.

To quantify the representational coupling between thalamic sub-region and DN subsystems, we concatenated the upper triangular vectors of their respective RSMs across all ToM events, and calculated the Pearson correlation between these concatenated vectors, followed by a Fisher’s Z transformation. We evaluated the significance of this coupling against a shuffled null distribution using an event-restricted permutation test with 1000 iterations. During each iteration, we applied a circular shift to the target ROI’s time series within each event window. By employing a substantial temporal offset (ranging from 10% to 90% of the event duration), we effectively disrupted the temporal alignment of representational content between regions while preserving the intrinsic autocorrelation and the block-diagonal structure of the RSM.

We defined the representational coupling effect as the difference between the true Fisher’s Z and the mean of the null distribution (ΔZ). At the network level, we derived one-tailed p-values based on the null distribution and applied FDR correction across all investigated thalamocortical pathways. We considered results significant at *q* < 0.05, indicating that the thalamus and DN subsystems share significant ToM-related information structures.

#### sEEG PPI analysis

We evaluated the directional communication between the thalamus and the DN using PPI analysis based on the normalized HFA time series described above, which were downsampled to 10 Hz. For each valid site pair, we constructed a GLM to estimate task-dependent changes in functional connectivity during mentalizing:

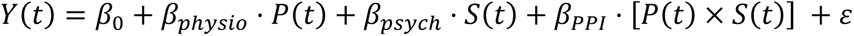

Here, *Y*(*t*) represents the HFA time series of the target site, and *P*(*t*) represents the HFA of the seed site (physiological variable). The psychological variable *S*(*t*) was defined as a binary vector (1 for ToM events, 0 otherwise), which we demeaned prior to computing the interaction term. We calculated the PPI term as the element-wise product of the seed physiological activity and the task-related psychological vector. To ensure specific directional inference, we investigated both feedforward (thalamus-to-DN) and feedback (DN-to-thalamus) pathways.

To evaluate the significance of the PPI effects, we implemented a permutation-based approach with 1000 iterations. In each iteration, we applied a circular shift to the psychological variable *S*(*t*) to generate a null distribution for the task-modulated connectivity (*β_PPI_*). Alternatively, to evaluate the intrinsic physiological baseline connectivity (*β_p_*_ℎ*ysio*_), we applied the circular shift directly to the seed physiological variable *P*(*t*) to break its inherent temporal alignment with the target site. To effectively break the specific task-related coupling while preserving the inherent rhythmic properties and autocorrelation of the neural signals, we enforced a minimum circular shift of 10 seconds and dynamically optimized the maximum temporal overlap between the shifted and original ToM events. Specifically, we initiated this overlap limit at 5% and incrementally relaxed it (in steps of 1%) until 1000 entirely unique time shifts (sampling without replacement) were successfully identified, with the final empirical overlap converging at approximately 16%. This adaptive constraint protocol mathematically ensures a sufficiently large pool of valid permutations while making surrogate psychological variables represent a distinct cognitive state.

At the group level, we pooled the estimated *β_PPI_* values across all site pairs for each specific thalamocortical pathway. We determined the significance by comparing these observed regional means against the permuted null distributions of means, followed by FDR correction across all tested pathways with significance set at *q* < 0.05.

#### PAC analysis

We quantified the dynamic coordination between low-frequency phase and high-frequency amplitude using cross-frequency phase-amplitude coupling (PAC). Using the continuous, broadband sEEG signal, we first applied a global Z-score normalization across the entire movie duration for each recorded site to standardize voltage variances. We then utilized the *pactools* package ^79^ to compute Tort’s Modulation Index (MI) for each site pair ^80^. This metric quantifies coupling by assessing the Kullback-Leibler divergence between the empirical amplitude distribution across phase bins and a uniform distribution. We calculated the MI across a frequency-frequency space, defining the low-frequency phase from 2 to 25 Hz in 1 Hz steps, and the high-frequency amplitude from 70 to 200 Hz in 6 Hz steps.

To systematically map the directionality of thalamocortical interactions, we evaluated three distinct coupling architectures: forward coupling (thalamic phase modulating DN amplitude), backward coupling (DN phase modulating thalamic amplitude), and local coupling (phase modulating amplitude within the same regional ROI). For each site pair and interaction type, we concatenated the data from all seven ToM events into a single continuous time series to compute the PAC. We then compared this to a baseline generated by bootstrapping 100 random continuous segments of identical total length sampled from the non-ToM periods. To account for potential global signal fluctuations and the aperiodic power bias, we derived a normalized PAC contrast ratio, defined as (PAC_ToM_ - PAC_Baseline_) / (PAC_ToM_ + PAC_Baseline_), yielding a metric isolated to task-specific coupling changes. To evaluate the significance of these interactions at the group level, we pooled all site pairs for each specific ROI-to-ROI pathways (e.g., DA thalamus to DN-B). We subjected the resulting PAC ratio maps to a one-sample cluster-based permutation test (1000 iterations) against zero and derived one-tailed p-values from these cluster-level statistics to identify significant PAC enhancements during mentalizing.

#### Thalamocortical PAC-gating analysis

To investigate whether thalamocortical PAC gates local activation and representation quality in the DN, we evaluated the correlation between the strength pairwise thalamic PAC and the functional relevance of their targeted DN sites.

We first quantified the local task-evoked activation for each DN site by averaging its normalized HFA across all ToM events. Next, we assessed each site’s contribution to the network representational geometry using a “virtual lesion” approach based on RSA (see “sEEG RSA analysis” section). We constructed a full network RSM across all ToM events using the HFA time series from all available sites within a specific cortical subsystem. For each target site, we then generated a lesioned RSM by excluding that specific site from the spatial pattern. We quantified the site’s representational contribution by calculating the Pearson correlation (*r_sim_*) between the upper triangular vectors of the full and lesioned RSMs. The final RSA contribution score was defined as 1 − *r_sim_*, where higher values indicate that the site carries critical multivariate information necessary for maintaining the network’s overarching representational structure.

To quantify the influence of thalamocortical PAC on these cortical sites, we computed the pairwise PAC index (between the 3-10 Hz phase of thalamic seed site and the 70-170 Hz amplitude of its corresponding DN target site) for every valid thalamocortical connection within a given subregion-network pair. To ensure the robustness of our correlation analyses, we implemented an automated outlier rejection procedure where any electrode pair exceeding four standard deviations from the mean across any of the neural metrics was excluded. Finally, we evaluated the modulatory role of thalamocortical PAC by calculating the one-tailed Pearson correlation across all valid thalamocortical electrode pairs between their pairwise PAC and the respective HFA activation, as well as the RSA contribution scores, of their cortical targets. Statistical significance was assessed using a permutation test with 1000 iterations. For each permutation, the cortical metrics (HFA or RSA contribution) were randomly shuffled across all pairs to construct a null distribution of correlation coefficients. Multiple comparisons across the four thalamic subregion within each cortical network were corrected using the FDR methods (*q* < 0.05).

### sEEG-fMRI temporal correlation analysis

To investigate the neurophysiological basis of the modality-specific differences observed in our RSA and to evaluate potential neurovascular decoupling within the thalamocortical circuit, we conducted a direct, electrode-to-voxel temporal correlation analysis between sEEG and fMRI signals.

For each sEEG electrode, HFA was extracted and cleaned following the exact pipeline detailed in the “HFA analysis” section. To accommodate the continuous cross-modal alignment, the smoothed and normalized HFA time series were downsampled to 10 Hz across the entire movie duration.

For fMRI BOLD signals, to optimize spatial specificity and avoid interpolation-induced blurring, signal extraction was performed in each participant’s native functional space. The MNI coordinates of all validated sEEG electrodes were mapped back to the individual native spaces using the inverse of the non-linear SyN warps and affine transformations generated via ANTs. Native BOLD time series were then extracted from a 4-mm radius spherical ROI centered on each transformed coordinate and subsequently normalized via Z-scoring.

To temporally align the electrophysiological signals with the BOLD responses, the sEEG HFA time series were convolved with a canonical double-gamma HRF. The HRF-convolved signals were then resampled to match the fMRI temporal resolution (TR = 1.72 s) via cubic spline interpolation.

We computed Pearson correlation coefficients between the HRF-convolved sEEG HFA and the native BOLD signals, which were subsequently Fisher Z-transformed. At the network level, the coefficients were first averaged across all valid electrodes within a given region for each fMRI subject, and these resulting subject-level means were subsequently tested against zero using one-sample t-tests against zero (two-tailed) across fMRI participants.

## Author contributions

Conceptualization: R.N.S and R.L. Data collection: E.D., J.M.S., D.L., Z.L. and J.D. Data analysis: R.L. and E.D., Writing – Original draft: R.L., Writing – Review & Editing: R.L., E.D., J.M.S., D.L., Z.L., P.K., J.D., X.J.C., B.C.B., G.R.T., R.N.S. Supervision and funding: R.N.S.

## Acknowledgements

The intracranial sEEG data were collected and generously shared by the Laboratory of Behavioral and Cognitive Neuroscience, led by Josef Parvizi at Stanford University, and supported by US National Institute of Neurological Disorders and Stroke R01NS137650 and R01NS078396, US National Institute of Mental Health R01MH134990 and R01MH109954. This project was supported in part by Canadian Institutes of Health Research (CIHR) and Natural Sciences and Engineering Research Council (NSERC) of Canada grants to R.N.S., a Fonds de Recherche du Quebec – Santé research scholar. X.J.C acknowledges the Canada Research Chairs program, CIHR and NSERC. B.C.B is supported by CIHR, NSERC, the Canada Research Chairs Program, Healthy Brains and Healthy Lives, and the Centre of Excellence in Epilepsy at the Neuro (CEEN). R.L. acknowledges postdoctoral scholar supported by CIHR (200883).

## Supplementary

**Figure S1.**
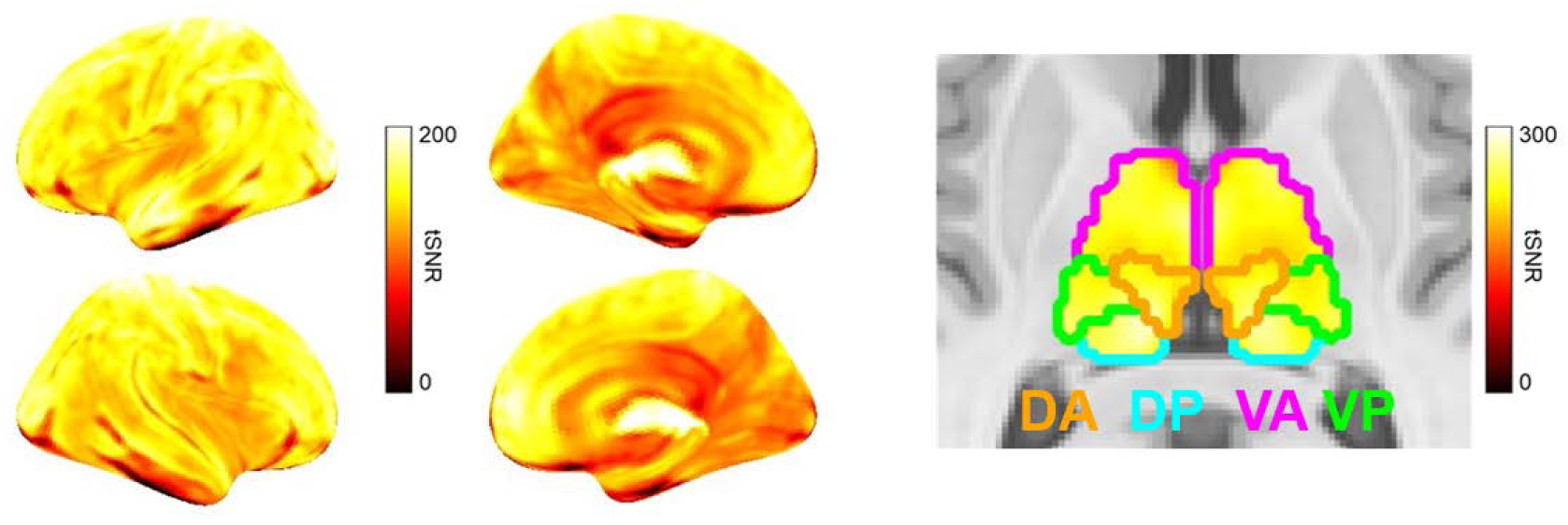
Temporal signal-to-noise ratio (tSNR) maps of the 7T multi-echo fMRI data. Group-averaged tSNR maps demonstrate robust signal quality across both cortical and subcortical regions.

**Figure S2.**
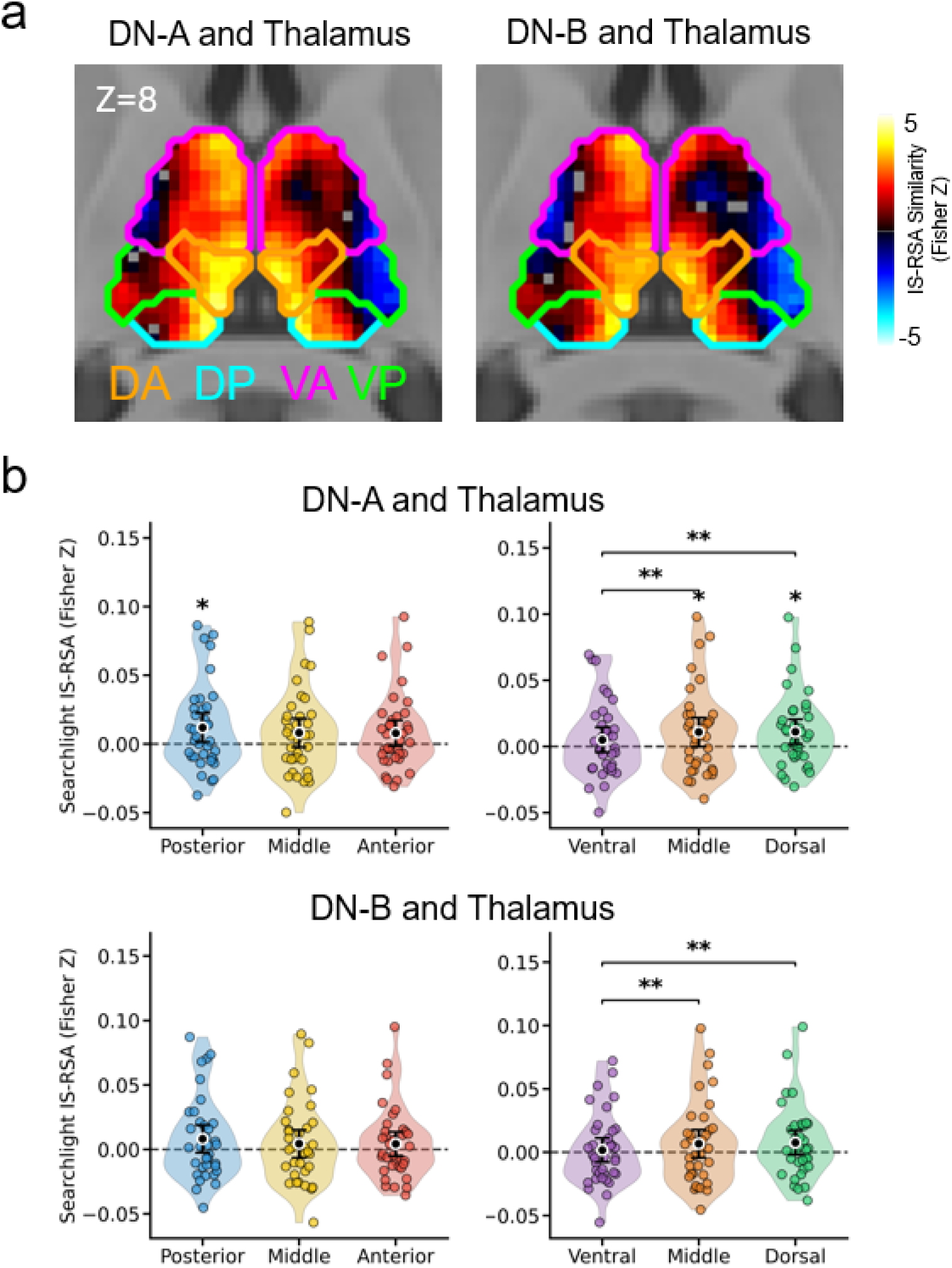
Sparse-sampling IS-RSA analysis. (a) Spatial topography of thalamocortical representational coupling revealed by searchlight sparse-sampling IS-RSA using a leave-one-out (LOO) framework. (b) Spatial binning analysis of the searchlight sparse-sampling IS-RSA along the anterior-posterior and dorsal-ventral axes. Each dot represents an individual subject, and error bars represent 95% confidence intervals. Pairwise comparisons among the three spatial bins were evaluated using two-tailed paired t-tests, while single-sample tests against zero (including the regional bin-specific tests and all ROI-level inferences) followed a directional one-tailed approach. * p < 0.05, ** p < 0.01 (FDR-corrected).

**Figure S3.**
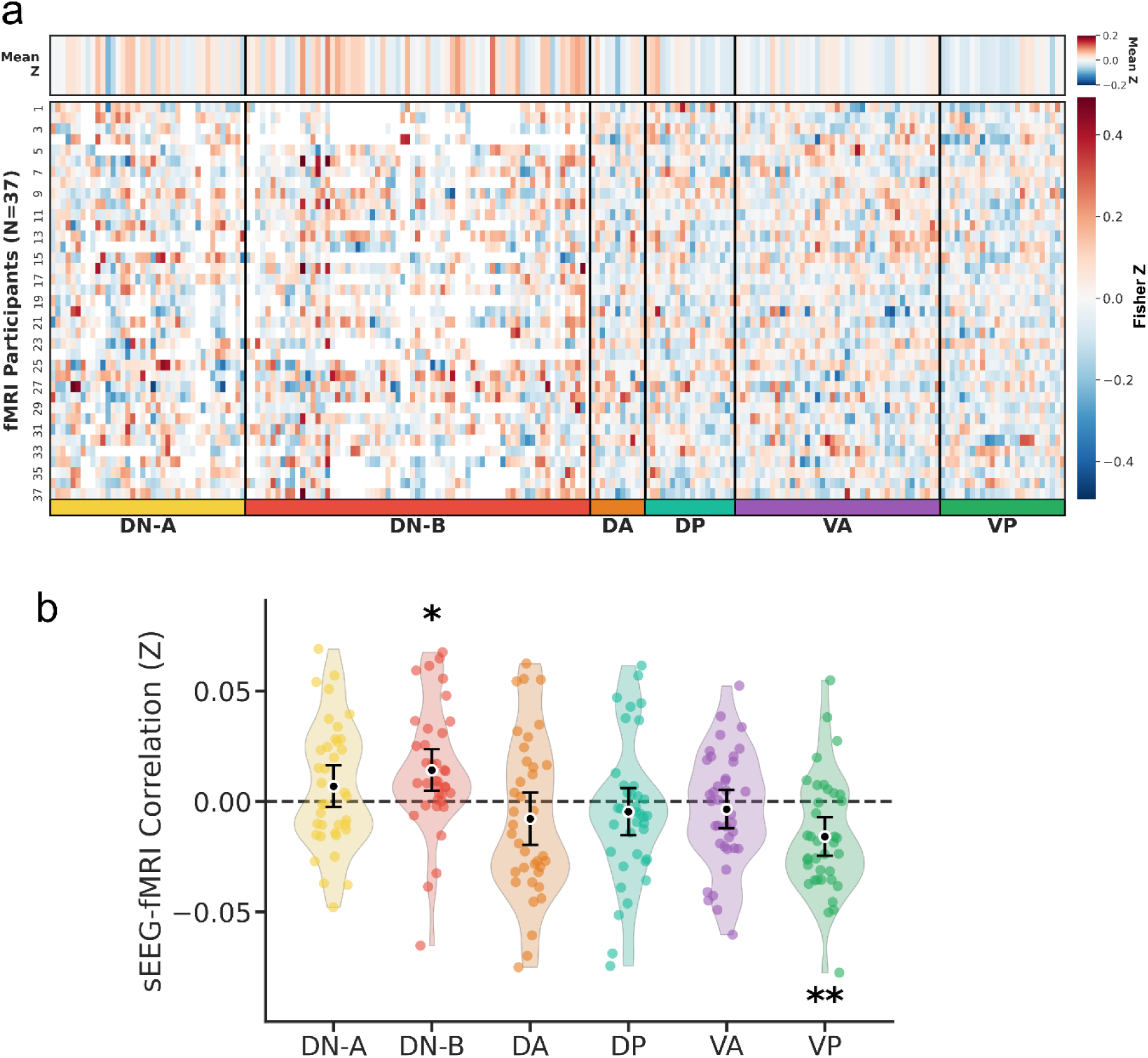
Cross-modal temporal correlation between sEEG and fMRI BOLD responses. (a) Brain-wide, electrode-level correlation matrix. The main heatmap displays the temporal correlation between the HRF-convolved sEEG HFA and the native-space fMRI BOLD signals across 37 independent fMRI participants (y-axis) and all valid sEEG recording sites (x-axis). Electrodes are grouped by cortical DN subsystems and thalamic ROIs. Color intensities reflect Fisher Z-transformed Pearson correlation coefficients. The top panel illustrates the mean Fisher Z across the fMRI cohort for each respective electrode. Notably, white spaces (NaNs) within the cortical DN regions denote electrodes where valid BOLD signals could not be extracted due to technical reasons (e.g., signal dropout at air-tissue interfaces or edge effects during nonlinear co-registration to individual native functional spaces). (b) Network-level average of the cross-modal temporal correlations. For each fMRI participant, Fisher Z values were first averaged across all valid electrodes within a given ROI to derive a subject-level mean. Each dot represents an individual fMRI participant. Error bars denote 95% confidence intervals. Statistical significance was evaluated across the fMRI cohort using one-sample t-tests against zero (two-tailed). * p < 0.05, ** p < 0.01 (FDR-corrected).

**Figure S4.**
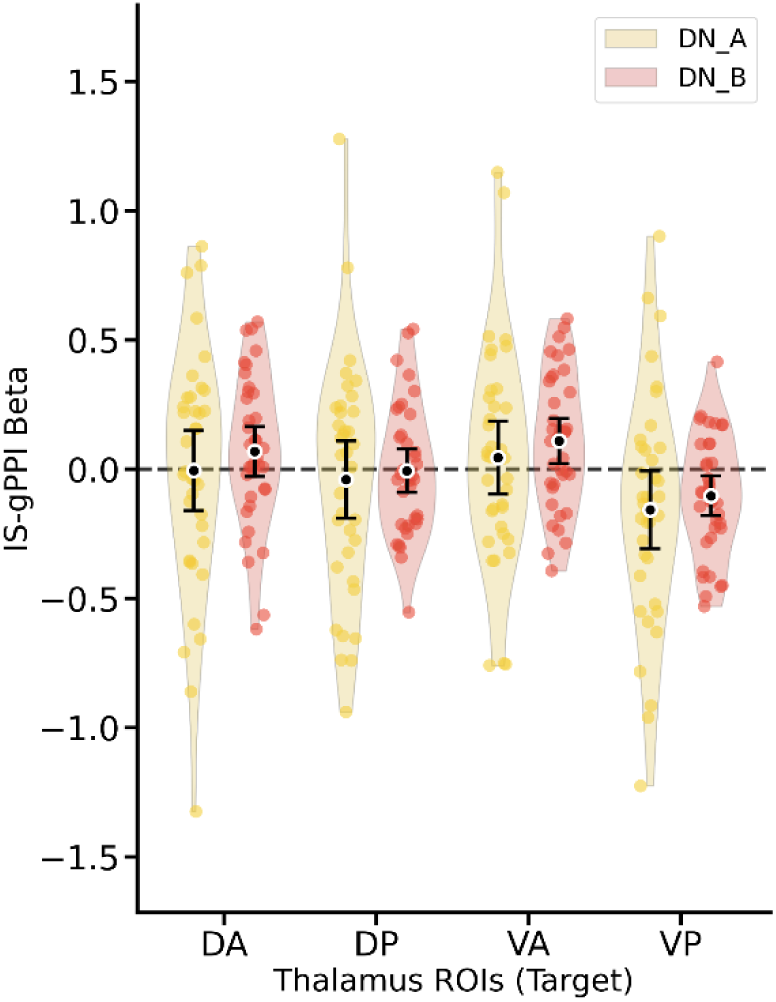
ROI-level fMRI IS-PPI results for corticothalamic (DN-to-Thalamus) connectivity during ToM. Violin plots display the task-modulated connectivity (IS-gPPI Beta) from default network subsystems to the four predefined thalamic ROIs. Error bars represent 95% confidence intervals, and each dot represents an individual subject.

**Figure S5.**
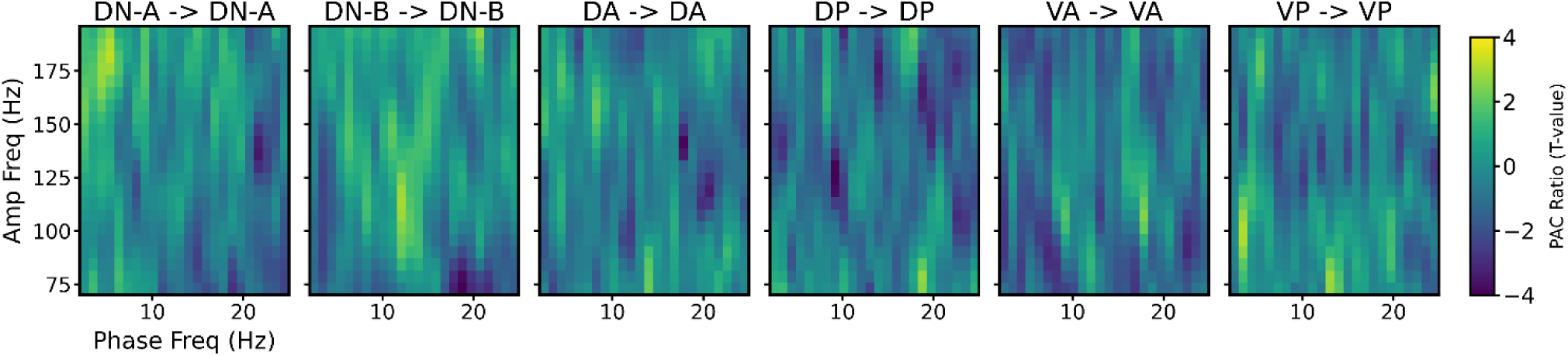
Local PAC within the DN and thalamic ROIs. ToM-related cross-frequency phase-amplitude coupling (PAC) from each ROI’s phase to its own high-frequency amplitude. Color maps represent the group-level normalized PAC contrast ratio (comparing ToM events against non-ToM baseline periods, expressed as T-values).

**Table S1.** Timeline and descriptions of ToM events in the animated short film *Partly Cloudy* (Adapted from Richardson et al., 2018)

| Event | Time | Duration (s) | Rephrased Description (Focusing on Mental States) |
| --- | --- | --- | --- |
| T1 | 4:00-4:10 | 10 | Fleeing from the baby shark, Peck lands on a happier cloud. Gus observes them, inferring that they are mocking him together. |
| T2 | 2:46-3:02 | 16 | Peck enviously watches another cloud create puppies. Gus misinterprets this as abandonment and looks anxious, causing Peck to feel embarrassed when caught staring. |
| T3 | 1:14-1:20 | 6 | A crying baby experiences a shift in emotional state, becoming happy and comforted after receiving a helmet. |
| T4 | 4:42-4:56 | 14 | Peck returns wearing football gear to resolve Gus's misunderstanding. He demonstrates that he didn't abandon Gus, but merely needed protective armor to safely transport Gus's hazardous creations. |
| T5 | 1:28-1:36 | 8 | The scene shifts to Gus, visually depicting his feelings of loneliness and isolation compared to the other cheerful clouds. |
| T6 | 3:48-3:58 | 10 | Startled by the dangerous baby shark Gus just created, Peck's attention is drawn to a neighboring cloud safely making chicks, prompting a contrast in expectations. |
| T7 | 1:50-1:54 | 4 | Peck and Gus share a joyful reunion, mutually expressing happiness and reaffirming their bond of friendship. |

